# Pollen dispensing schedules in buzz-pollinated plants: Experimental comparison of species with contrasting floral morphologies

**DOI:** 10.1101/2020.08.04.235739

**Authors:** Jurene E. Kemp, Mario Vallejo-Marín

## Abstract

1. In buzz-pollinated plants, bees apply vibrations to remove pollen from anthers that have small apical pores or slits. These poricidal anthers potentially function as mechanism to stagger pollen release, but this has rarely been tested across plant species differing in anther morphology.
2. In *Solanum* section *Androceras*, three pairs of buzz-pollinated *Solanum* species have undergone independent evolutionary shifts between large- and small-flowered taxa. These shifts in flower size are accompanied by replicate changes in anther morphology, and we used these shifts in anther morphology to characterise the association between anther morphology and pollen dispensing schedules. We characterised pollen dispensing schedules by applying simulated bee-like vibrations directly to anthers to elicit pollen release. We then compared pollen dispensing schedules across anther morphologies, and we further investigated how vibration velocity affects pollen release. Finally, we assessed whether particular anther traits, presented in the Buchmann-Hurley model, can predict pollen dispensing schedules.
3. We show that replicate transitions in *Solanum* anther morphology are associated with consistent changes in pollen dispensing schedules. We found that small-flowered taxa with small anthers release their pollen at higher rates than their large-flowered counterparts, showing an association between general anther morphology and pollen dispensing. Further, higher vibration velocities resulted in quicker pollen dispensing and more total pollen released, which suggested that bees that produce high-energy vibrations can access more reward than bees producing low-energy vibrations. Finally, both the pollen dispensing rate and the amount of pollen released in the first vibration were negatively related to anther wall area, but, surprisingly, we did not observe any association between pore size and pollen dispensing.
4. Our results provide the first empirical demonstration that the pollen dispensing properties of poricidal anthers depend on both floral characteristics and bee vibration properties, and suggest that morphological modification of anthers could provide a mechanism to exploit different pollination environments.

## Introduction

Most flowering plant species rely on animals to transport pollen between conspecific flowers for fertilisation (Ollerton *et al.*, 2011). Despite the widespread reliance on animals as pollen vectors, animal pollination can limit pollen dispersal, particularly when visitors actively collect pollen, making it unavailable for fertilisation (Harder & Wilson, 1994; Minnaar *et al.*, 2019). In nectarless plants, where pollen serves both pollinator reward and vehicles for male gametes, we expect selection to favour adaptations that limit the fitness costs associated with pollen consumption, whilst releasing enough pollen to ensure sufficient pollinator visits (Harder & Thomson, 1989; Harder & Barclay, 1994; LeBuhn & Holsinger, 1998; Vallejo-Marín *et al.*, 2009).

Plants can theoretically mitigate the fitness costs associated with pollen consumption by optimizing their pollen dispensing schedules (i.e., the rate at which pollen is released across visits) to the visitation rates and grooming behaviours of pollinators (Harder & Thomson, 1989; Harder & Wilson, 1994; LeBuhn & Holsinger, 1998). Theoretical models predict that pollen should be gradually released across multiple visits when pollinator visits are unlimited and when their grooming behaviours result in high diminishing fitness returns (i.e., when the total amount of pollen transferred decreases with the amount of pollen collected per visit) (Harder & Thomson, 1989; Harder & Wilson, 1994). In contrast, when pollinator visits are limited and have high per-visit transfer efficiencies, pollen should be released across fewer visits (Harder & Thomson, 1989; Harder & Wilson, 1994). Thus, when pollinators collect pollen and exhibit low per-visit transfer efficiencies, as in many bee pollinated taxa, models predict that selection will favour pollen being gradually released across multiple visits if pollinators are abundant (Thomson, 1986; Harder & Thomson, 1989; Holsinger & Thomson, 1994; Schlindwein *et al.*, 2005; Harder & Johnson, 2009). However, restricting pollen removal excessively might be in conflict with the requirements of pollinators that collect pollen, and pollinators might avoid visiting flowers that release too little pollen per visit (Harder, 1990a). Pollen dispensing schedules thus represent the evolutionary outcome of selective pressures acting on plants and mediated by pollinators, and are particularly important in plants that offer pollen as the primary reward (Harder & Barclay, 1994).

Plants can dispense pollen gradually through moderating anther maturation within a flower or by staggering flower opening within a plant individual (Sargent, 2003; Castellanos *et al.*, 2006a; Li *et al.*, 2014). Another potential solution to limit per-visit pollen collection is to physically restrict access to pollen, as is done in poricidal anthers that contain pollen inside the anthers (Harder & Barclay, 1994; Dellinger *et al.*, 2019c). Poricidal anthers are associated with buzz-pollination (Buchmann, 1983). During buzz pollination, bees vibrate flowers using their thoracic muscles. The vibrations transmitted through the anthers cause pollen to be ejected through the apical pores (De Luca & Vallejo-Marín, 2013; Vallejo-Marín, 2019). Buzz-pollinated flowers are typically nectarless and solely offer pollen as reward (Buchmann, 1983; Faegri, 1986; Vallejo-Marín *et al.*, 2010), which makes them a likely candidate for gradual pollen release. Buzz-pollination is widespread among plants, with more than 20,000 species possessing poricidal anthers (Buchmann, 1983), and about half of bee species capable of producing floral vibrations (Buchmann, 1983; Cardinal *et al.*, 2018). The floral and anther morphology of buzz-pollinated flowers is very diverse, and even plants with poricidal anthers vary widely in their stamen morphology, both between and within species (Buchmann, 1983; Dulberger *et al.*, 1994; Corbet & Huang, 2014; Vallejo-Marín *et al.*, 2014; Dellinger *et al.*, 2019a,b). Similarly, the vibration properties of bees vary across and within species (King & Buchmann, 2003; Arceo-Gómez *et al.*, 2011; Corbet & Huang, 2014; Arroyo-Correa *et al.*, 2019; De Luca *et al.*, 2019; Pritchard & Vallejo-Marín, 2020), and previous empirical work demonstrates that vibration properties, particularly their amplitude, affect pollen release (Harder & Barclay, 1994; King & Buchmann, 1996; De Luca *et al.* 2013). However, we know relatively little about how plant species with different anther and floral morphologies vary in their pollen release schedules (but see Harder & Barclay 1994, Dellinger *et al.* 2019), including their response to vibrations of different amplitudes.

Anther properties, such as shape, length, natural frequency and pore size, are theoretically expected to influence pollen release (Buchmann & Hurley, 1978; Vallejo-Marín 2019). In what remains the only biophysical model of buzz pollination, Buchmann & Hurley (1978), modelled a poricidal anther with a simple geometry (a rectangular box) and an anther pore at one end. The anther vibrates along a single spatial axis perpendicular to the longest anther dimension, with pollen grains bouncing against the interior walls of the anther locule. In this model, pollen release occurs when the kinetic energy of the vibrating anthers is transferred to the pollen grains, which are then ejected through the pore. The higher the energy, the higher the pollen release rate. Pollen grains can gain energy as they bounce against the interior of the anther walls or against each other. Thus, the internal surface area of the anther locule along the axis of vibration (*A*) is positively related with the rate of change in energy of pollen grains (equation 9 in Buchman & Hurley 1978). However, energy can also be lost as pollen grains escape the anther through the pore, and therefore the area of the anther pore (*A’*) is negatively related to changes in pollen energy (eqn. 9). Because *A’* is positively related to the rate at which pollen grains are released from anthers (equation 10), but negatively related to changes in pollen’s kinetic energy, the relationship between anther pore size and pollen release dynamics is not immediately obvious, as dynamics change as the anther empties while vibrated. Numerical integration of Buchman & Hurley’s model suggests that both increased anther wall area and pore size should result in faster times to empty anthers, and larger anther volumes should result in slower pollen release (B. Travacca & M. Vallejo-Marin, unpublished), but clearly more theoretical work is needed in this area.

In contrast to the absence of work on associations between anther morphology and pollen release rates, the effect of the type of vibration applied (e.g., vibration frequency, amplitude and duration) on pollen release has received more attention (Harder & Barclay, 1994; De Luca *et al.*, 2013; Rosi-Denadai *et al.*, 2018). A study using bee-like vibrations (in terms of frequency, duration and amplitude) showed that pollen release is more strongly affected by amplitude and duration than by the frequency of the vibration (De Luca *et al.*, 2013). However, previous studies assessing the effects of vibration properties on pollen release have focused on the amount of pollen released during a single vibration, and thus it has not been possible to build a full picture of pollen release curves following multiple consecutive buzzes.

Here, we characterise pollen dispensing schedules across three pairs of closely related buzz-pollinated taxa in the genus *Solanum* section *Androceras* (Solanaceae). These six taxa represent three pairs of independent evolutionary transitions from large flowers with large anthers and small pores, to small flowers with small anthers and large pores (Whalen 1978; 1979; Vallejo-Marin *et al.*, 2014; Rubini-Pisano *et al*., in prep). These large-to small-flower transitions bear the classic hallmark traits of shifts in mating system from outcrossing towards increased self-pollination (Vallejo-Marín *et al.*, 2014; Rubini-Pisano *et al.*, in prep), but have also been suggested to be associated with pollinator shifts (Whalen, 1978). Regardless of the cause of the shift in flower and anther morphology, these changes allow us to investigate the association between anther morphology and pollen dispensing schedules in closely-related taxa. We first investigate whether these replicate transitions in general floral morphology are associated with replicate changes in pollen dispensing schedules. We then use one of these evolutionary transitions from large-to small-flowered species to investigate the extent to which pollen dispensing schedules depend on vibration velocity. Finally, we test whether the parameters in the Buchmann & Hurley (1978) model can predict pollen release rates in the six focal taxa. The replicate evolutionary transitions in anther morphology in three closely-related clades of *Solanum* provides an ideal opportunity to establish how pollen dispensing schedules of buzz-pollinated plants vary with anther morphology, and how this is influenced by vibration properties.

## Materials and Methods

### Study system

The genus *Solanum* is often used as a model system for studying buzz-pollination (Vallejo-Marín 2019). *Solanum* contains c. 1,350 species with nectarless flowers, representing about half of the species diversity in the family Solanaceae (Särkinen *et al.*, 2013). *Solanum* flowers are pentamerous, usually radially symmetric, and bear poricidal anthers (Harris, 1905; Knapp, 2002). During buzz-pollination, bees typically grab the base of the anthers using their mandibles whilst curling the ventral side of their bodies around the anthers. They then use their indirect flight muscles to vibrate the anthers to instigate pollen release (King *et al.*, 1996). *Solanum* section *Androceras* is a monophyletic clade consisting of approximately 12 annual and perennial species distributed in Mexico and the southern USA (Whalen, 1978; Stern *et al.*, 2010). All taxa in section *Androceras* are heterantherous, i.e., have two or more morphologically differentiated sets of anthers in the same flower. In heterantherous species, one set of anthers is usually associated with pollinator attraction and reward (“feeding” anthers), while the other set contributes disproportionately to fertilising ovules (“pollinating” anthers) (Vallejo-Marín *et al.*, 2009). The section is divided into three series: *Androceras*, *Pacifum*, and *Violaceiflorum* (Whalen, 1979). Parallel shifts in flower and anther morphology have occurred within each of these three series (Vallejo-Marín *et al.*, 2014) (Figs. 1 & S1). In series *Androceras*, *S. rostratum* has large flowers, and long anthers with small anther pores, whereas *S. fructu-tecto* has small flowers with small anthers and large anther pores (referred to as the SR-SF clade, hereafter). Similar morphological shifts are present in series *Pacificum* for *S. grayi* var. *grandiflorum* (large flowers) and *S. grayi* var. *grayi* (small flowers) (referred to as the SGN-SGG clade, hereafter), and in series *Violaceiflorum* for *S. citrullifolium* (large flowers) and *S. heterodoxum* (small flowers) (referred to as the SC-SH clade hereafter). These shifts in floral morphology is potentially associated with shifts in mating system (i.e., from outcrossing to selfing), although this has not been empirically confirmed (Vallejo-Marín *et al*., 2014, Rubini-Pisano *et al*., in prep.). Within these species pairs, the pollinating and feeding anthers are less differentiated in the small flower types than in the large flower types (Vallejo-Marín *et al.*, 2014). We refer to these two distinct floral morphologies as the ‘small flower type’ and ‘large flower type’ throughout, following Rubini-Pisano *et al.* (in review).

**Figure 1.**
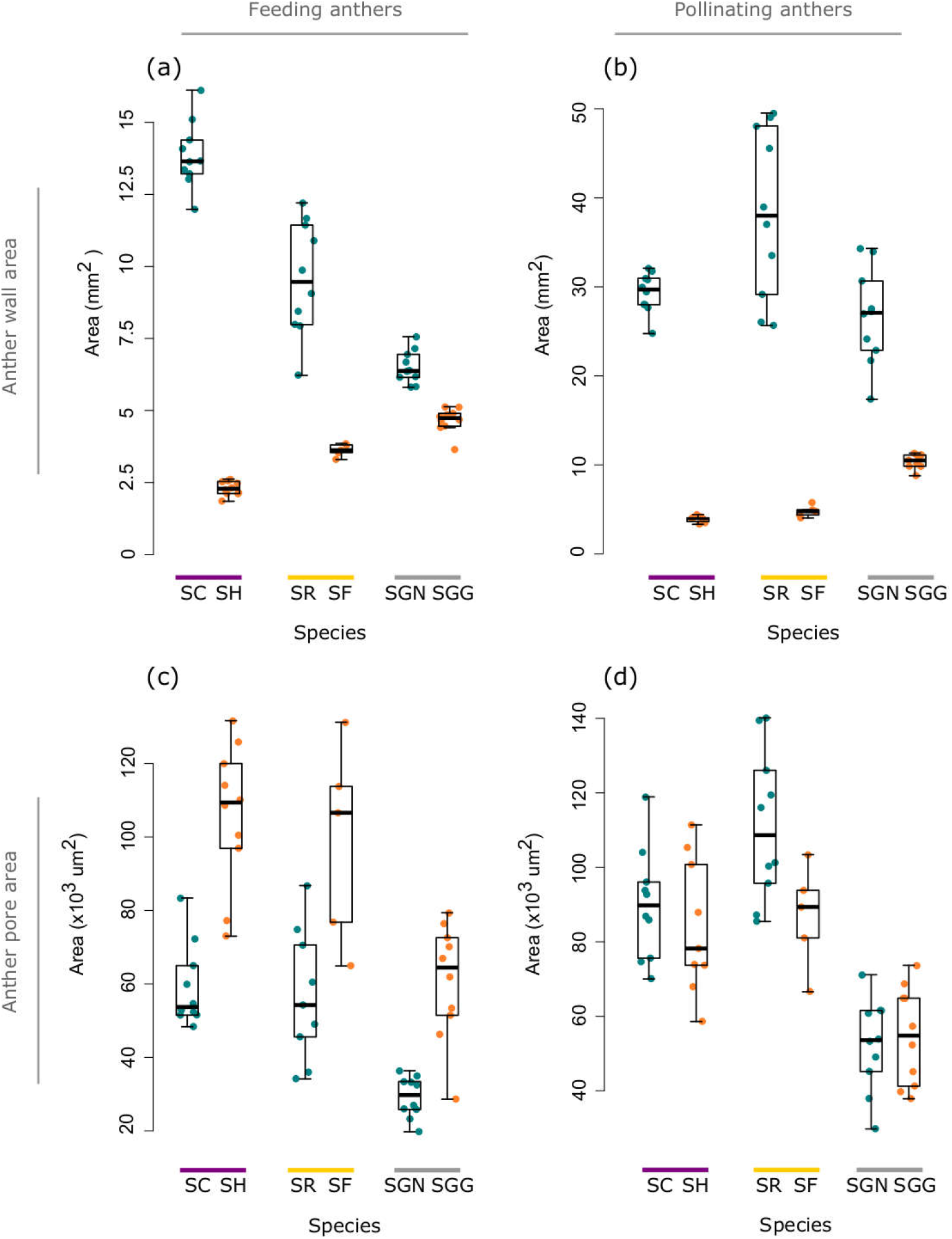
Anther wall (top panel) and anther pore areas (bottom panel) for the feeding (lefthand panel) and pollinating (righthand panel) anthers of three pairs of taxa in *Solanum* section *Androceras* (Solanaceae). Anther wall area was calculated following Buchmann & Hurley (1978) as the product of the length and breadth of anthers. Anther pore area was measured from SEM photographs. We show the values of individual anthers, and not the summed values used in the analyses. The six studied taxa belong to three phylogenetic clades indicated by the purple, yellow and grey lines above taxon names. Within each clade, blue dots indicate the large flowered type and orange dots indicate the small flowered type. Species names are: SC =*Solanum citrullifolium*; SH = *S. heterodoxum*; SR = *rostratum*; SF = *S. fructu-tecto*; SGN = *S. grayi* var. *grandiflorum*; SGG = *S. grayi* var. *grayi.*

### Seed collection and plant growth

Seeds were collected from natural populations in Mexico between 2007 and 2010, except for *S. citrullifolium* which was obtained from the Solanaceae collection at Radboud University, Netherlands (Experimental Garden and Genebank Solanaceae collection). All of these natural populations occurred within a 150 km radius, and these areas have similar water vapour pressures (Worldclim 2, Fick & Hijmans, 2017, Table S1). For each plant species, we used seeds from a single population. Accession numbers and sample localities are provided in Table S1. Germination was induced by treating seeds for 24h with 1000 ppm aqueous solution of gibberellic acid (GA3; Sigma-Aldrich, Dorset, UK), following the method of Vallejo-Marín *et al.* (2014). Two to four weeks after germination, seedlings were transplanted to 1.5 L pots and kept in a pollinator-proof greenhouse with natural light supplemented with compact fluorescent lamps to provide at least 16hours of daylight. Temperatures were kept between 16 and 25°C (day and night).

### Synthesising bee-like vibrations and playback system

To characterise pollen dispensing schedules, we applied mechanical vibrations to flowers. We used Audacity v2.1.3 (http://audacity.source-forge.net/) to generate an artificial vibration (stimulus) with similar properties (amplitude and frequency) to the vibrations that bees produce when extracting pollen (De Luca & Vallejo-Marín, 2013). The vibrations consisted of a pure tone (300 Hz) sine wave made of five consecutive pulses of 200 milliseconds (ms) each with 200 ms of silence between pulses (i.e., total stimulus length = 2 seconds). Each pulse had a fade-in feature of 10 ms (Fig. S2). Multiple short buzzes with a single dominant frequency characterise the floral vibrations of buzz pollinating bees (De Luca & Vallejo-Marín 2013, Pritchard & Vallejo-Marín, 2020), and a 300 Hz frequency was selected to capture the frequency of floral vibrations typically produced by medium-sized bees, including bumblebees (De Luca & Vallejo-Marín, 2013; Switzer & Combes, 2017; Arroyo-Correa *et al.*, 2019; De Luca *et al.*, 2019; Pritchard & Vallejo-Marín, 2019). Previous studies have shown that variation in frequency does not alter pollen release in a single vibration, but rather, higher accelerations result in higher pollen release (De Luca *et al.*, 2013). Theoretically, if flowers are vibrated at the natural frequency of anthers, higher pollen release could be induced due to the higher accelerations associated with resonance (King & Buchmann, 1996). For our six taxa, Nunes (2020) showed that only the feeding anthers of *S. grayi grayi* might resonate at 300 Hz (mean natural frequency = 294 Hz). We thus used a single ecologically relevant frequency for all experiments (i.e. 300 Hz), and we vary acceleration in the experiments (described below). The vibration amplitude was calibrated as described below to obtain the appropriate accelerations for each experiment.

The synthesised vibrations were applied using a custom-made vibration transducer system (A. Gordon and M. Vallejo-Marín, unpublished). This playback system consisted of a vibration transducer speaker (Adin S8BT 26W, Shenzhen, China) with a vibrating metal plate. We attached a metal rod (15 cm long with a 0.5 cm diameter) using an ethyl cyanoacrylate glue (Loctite UltraGel Control, Düsseldorf, Germany) and plastic supports at the base. We fixed a pair of featherlight forceps (D4045, Watkins & Doncaster, Leominster, UK) to the distal end of the metal rod at a 90° angle using a metal clip. We used laser vibrometry to calibrate the vibration amplitude and check that the playback frequency matched the input frequency. Briefly, we deployed a PDV-100 laser vibrometer (PDV-100, Polytec, Waldbronn, Germany) and focused the laser close to the tip of the forceps (~1 cm from the tip), where we had placed a small amount of reflective tape. The laser beam was aimed on the forceps perpendicular to the main axis of displacement of the transduction system. The vibration signal was played in Audacity using a laptop computer connected to the transduction system. The frequency and amplitude of the vibration was checked using VibSoft-20 data acquisition and software (Polytec, Waldbronn, Germany). Peak amplitude velocity of the vibration was adjusted using the volume control in the computer until the desired velocity was obtained. This calibration was done at the beginning of every day of the experiment, and again after pollen was extracted from 3-5 flowers.

### Pollen extraction and counting

Experimental flowers were brought from the glasshouse between 7h00 and 9h00 on the first day of flower opening in a closed container with wet floral foam (Oasis Floral Products, Washington, UK), to prevent flowers from drying out. Closed flower buds were tagged on the previous day in the late afternoon to ensure that only newly opened flowers are used in the experiments. All pollen extraction treatments were done within three hours after the flowers were picked. Maximum ten flowers were used per day, depending on availability, and multiple plant species were used each day. The artificial stimuli were applied to flowers in May 2019 at room temperature (22°C) in an indoor airconditioned laboratory at the University of Stirling (UK).

An individual flower, including the pedicel, was attached to the vibration transducer system using the forceps. The forceps were used to hold the flower at the base of the five anthers (cf. De Luca *et al*., 2013), and vibrations were thus directly transferred to the anthers in a similar manner as when bees vibrate flowers. A single vibration (consisting of five 200 ms buzzes as described above), was applied to the anthers of a flower, and the ejected pollen was collected in a 1.5 ml microcentrifuge vial. Each flower was subjected to 30 consecutive vibrations for a grand total of 30 s of buzzing time. Pollen was collected in separate vials for vibrations 1—10, 15, 20, 25, and 30. At the end of the trial, we removed the anthers of the flower and placed them in a centrifuge tube with 200 μl of 70% ethanol. The remaining pollen in the flower was extracted later with the help of a sonicating bath (D00351, Premier Farnell Ltd., Leeds, UK), which allowed us to calculate the total amount of pollen grains in each flower we used in our pollen dispensing trials.

To estimate the number of pollen grains in each sample (including samples of the pollen remaining in anthers), we used a particle counter (Multisizer 4e Coulter Counter, Indianapolis, USA). Each pollen sample (suspended in 200 μl 70% ethanol) was added to 20 ml 0.9% NaCl solution. For each sample, the amount of pollen was counted in two 1 ml subsamples. The pollen counts for these subsamples were averaged and multiplied by 20 to obtain the total pollen count. For samples with higher pollen concentrations, such as those containing the pollen remaining in the anther, we added the pollen samples to 100 or 200 ml NaCl solution and multiplied the averaged pollen count by 100 or 200 respectively to obtain the total pollen count. Blank samples, containing only 0.9% NaCl solution were run at the beginning of a session and regularly between samples to ensure calibration accuracy.

### Characterising pollen dispensing curves

To characterize the pollen dispensing curves, we fitted exponential decay curves for each flower using the *nls* function in R ver. 3.6.0 (R Core Team, 2019). The decay curves followed the function:

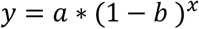

where *y* represents the percentage of pollen released in a vibration, and *x* represents the vibration number. The parameter *a* represents the intercept of the pollen dispensing curve, and the parameter *b* represents the percentage decrease in the amount of pollen released in each successive vibration (e.g., if b = 0.3, then 30% of the remaining pollen is released in each successive vibration). Model parameters were estimated separately for each flower using the *nls* function. The percentage pollen released per vibration was calculated by dividing the amount of pollen released per vibration by the total amount of pollen in a flower.

### Measuring anther traits

For each flower used in our trials, we measured three traits for the pollinating and feeding anthers separately, after pollen had been extracted. We measured: (1) the anther length, (2) the anther breadth at base of the anther, and (3) the area of the pores (Fig. S3). Anther length and breadth were measured using a dissection microscope and callipers. For each flower, all four feeding anthers were measured, and lengths and breadths were averaged across the feeding anthers to calculate a single value per flower. To calculate the anther pore area, we took SEM photographs of one feeding and one pollinating anther per flower, and we measured the area of the pores from photographs using ImageJ v1.52 (Schneider *et al.*, 2012). Variation in traits between species can be seen in Fig. 1.

### Variation in pollen dispensing curves across flower types

To assess whether pollen dispensing schedules vary between flower types, we subjected *Solanum rostratum, S. fructu-tecto, S. grayi* var. *grandiflorum, S. grayi* var. *grayi, S. citrullifolium*, and *S. heterodoxum* flowers to simulated vibration stimuli as described above. All vibration stimuli had a peak velocity of 80 mm/s. We used this velocity as it corresponds with the higher end of velocities which have been recorded for bees on flowers (De Luca & Vallejo-Marín, 2013). Pollen dispensing curves were characterized for ten flowers per species, except for *S. fructu-tecto* where only five flowers were used.

To compare the pollen dispensing curves, we extracted two response variables from each curve. Firstly, we extracted the amount of pollen released from the first simulated vibration, which reflects the amount of pollen reward a pollinator will receive in a single visit. Secondly, we estimated the rate of pollen release (i.e., the dispensing rate) as represented by *b* in the exponential decay function. For example, if *b* = 0.2, then 20% of the remaining pollen in the anther is released in each sequential vibration. After testing for normality, we used nonparametric Wilcoxon rank sum tests to compare the two response variables between the small and large flower types within each of the three phylogenetic clades separately. We chose to perform three separate within-clade tests, due to both the non-normality of our data and our small sample sizes. Because we performed multiple tests, we used the Holm method to adjust p-values. In addition to releasing pollen across multiple vibrations, plants can potentially further stagger pollen release across hours or days. We tested for this by comparing the total percentage of pollen that was released from anthers across the 30 vibrations between anther types within clades using Wilcoxon rank sum tests with Holm-adjusted p-values.

### Effect of vibration properties on pollen dispensing

We assessed the influence of vibration properties on pollen dispensing schedules by applying simulated vibrations to flowers of two species: *Solanum citrullifolium* (large flower type) and *S. heterodoxum* (small flower type). We focused on the effects of variation in velocity because previous work has shown that velocity is positively associated with the amount of pollen released in a single vibration (De Luca *et al.*, 2013). We applied three peak velocity treatments: 80, 40, and 20 mm/s. These values are within the range of previously recorded bee vibrations (De Luca *et al*., 2013). Dispensing curves were characterized for nine to ten flowers for each treatment and species (Table 1). We used the same response variables as in the previous section (i.e., percentage of pollen released in the first vibration and the dispensing rate), and we compared these response variables across the three velocity treatments within each species separately using analysis of variance (ANOVA) and Tukey posthoc tests. Further, we compared the total amount of pollen released per treatment within each species using ANOVA and Tukey posthoc tests.

**Table 1.**
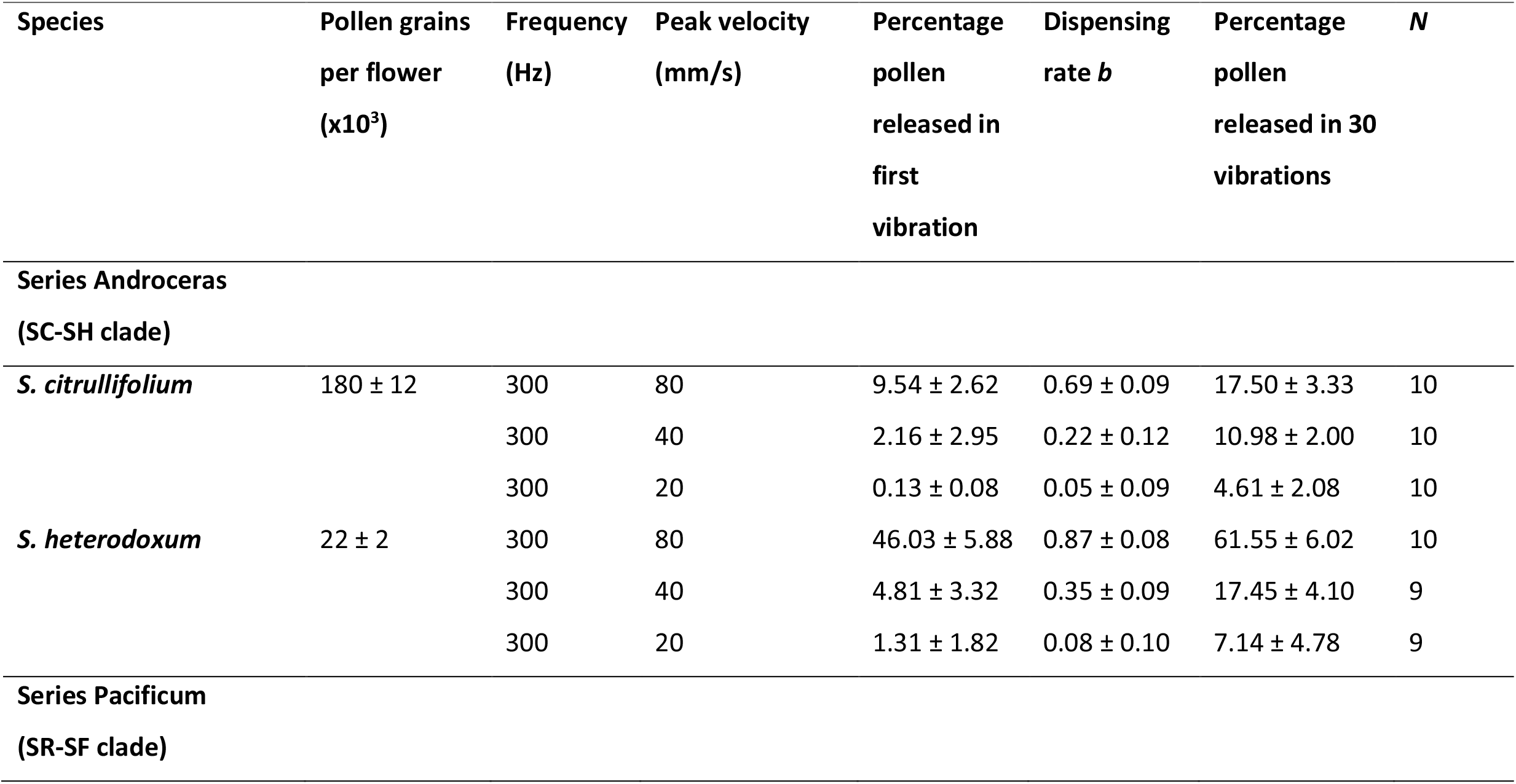
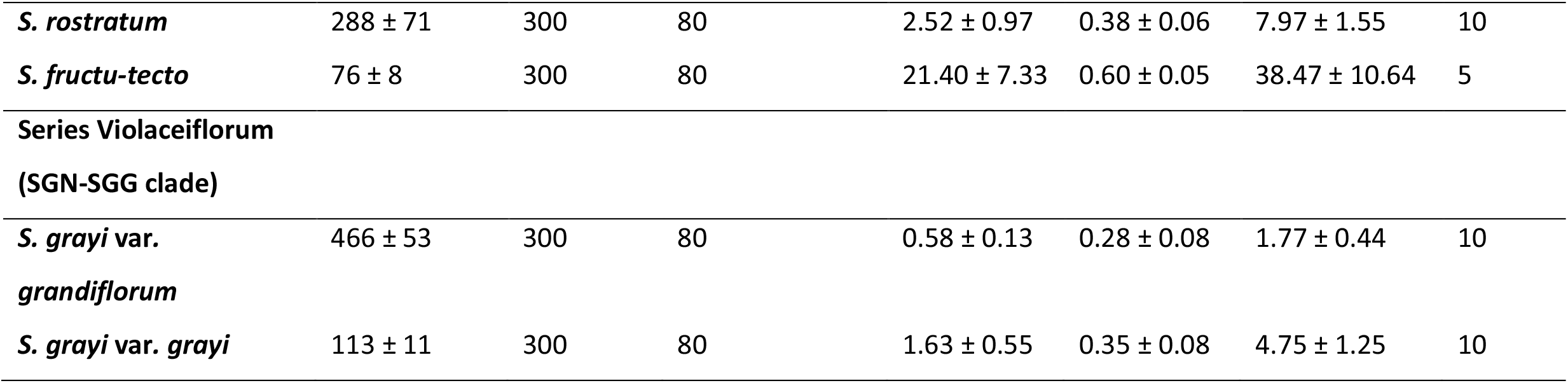
Pollen dispensing schedule metrics for six *Solanum* taxa during simulated buzz pollination.. In each treatment, flowers were subjected to 30 simulated vibrations. All species were subjected to vibrations with a peak velocity of 80 mm/s and frequency of 300 Hz (which equals an acceleration of 151 m/s2 and a displacement of 42 μm). Two species (*Solanum citrullifolium* and *S. heterodoxum*) were also exposed to two lower velocity vibrations, i.e., 40 mm/s (acceleration = 75 m/s^2^; displacement = 21 μm) and 20 mm/s (acceleration = 38m/s^2^; displacement = 11 um). Median ± standard error. *N* = sample size.

### Association between pollen dispensing schedules and anther traits

If consistent differences are found between large and small flowered taxa, the question of what causes those differences remains. One hypothesis is that the anther traits specified in the Buchmann & Hurley (1978) model predicts the rate of pollen release. The anther traits Buchmann & Hurley (1978) used in their model of pollen release from poricidal anthers include the area of the pore (*A’*), the area of the anther locule along the wall perpendicular to the movement direction of the anther (*A*), and the internal volume of the anther’s locule (ϑ). Because of the complexity of anther shapes and the technical difficulty of accurately estimating the internal dimensions of the anther locule, we used external anther area (length x breadth of the anther; Buchmann & Hurley, 1978; Figure S3) as a proxy of locule area *A*. We kept the pollinating and feeding anther traits separately, rather than combining their areas, because of the morphological differences between these anther types which may have different effects on pollen release. Further, because flowers contain four feeding anthers and one pollinating anther, we multiplied the values of both the pore area and the anther wall area of the feeding anther by four to obtain total areas for each flower. Because kinetic energy was kept constant in our pollen extraction trials, we could directly assess the association between pollen dispensing schedules and anther traits.

To determine whether the amount of pollen released in the first vibration is related to anther traits, we implemented a negative binomial mixed effect model (LMM) in *lme4* (Bates *et al.*, 2015). We used the amount of pollen released in the first vibration as response variable and the four anther traits (i.e., pore area and anther wall area for pollinating and feeding anthers separately) as predictor variables. Because the total amount of pollen grains varied between flowers, we also included the total amount of pollen grains per flower as offset in the model. We added species identity as a random effect. Similarly, we evaluated the association between pollen dispensing rate and anther traits. Because the dispensing rate metric (*b*) is bounded between 0 and 1, we conducted a logistic mixed effect model (GLMM) using the dispensing rate as the response variable and the four anther traits as the predictor variables, with species identity as a random effect.

## Results

### Pollen dispensing

Plant species varied in the amount of pollen grains that were present in flowers, with *Solanum heterodoxum* containing the fewest pollen grains (22 × 10^3^ ± 8 × 10^3^; median ± se) and *S. grayi* var. *grandiflorum* containing the most (446 × 10^3^ ± 17 × 10^3^; Table 1). Small flowered taxa consistently contained fewer pollen grains than their large flowered counterparts. The amount of pollen released decreased gradually with an increasing number of vibrations applied, and no pollen was released by the 30^th^ vibration (Fig. 2), despite large amounts of pollen remaining in the anthers (Fig. 3c). The pollen dispensing curves thus showed exponential decay, with the majority of pollen released in the first vibration (Fig. 2).

**Figure 2.**
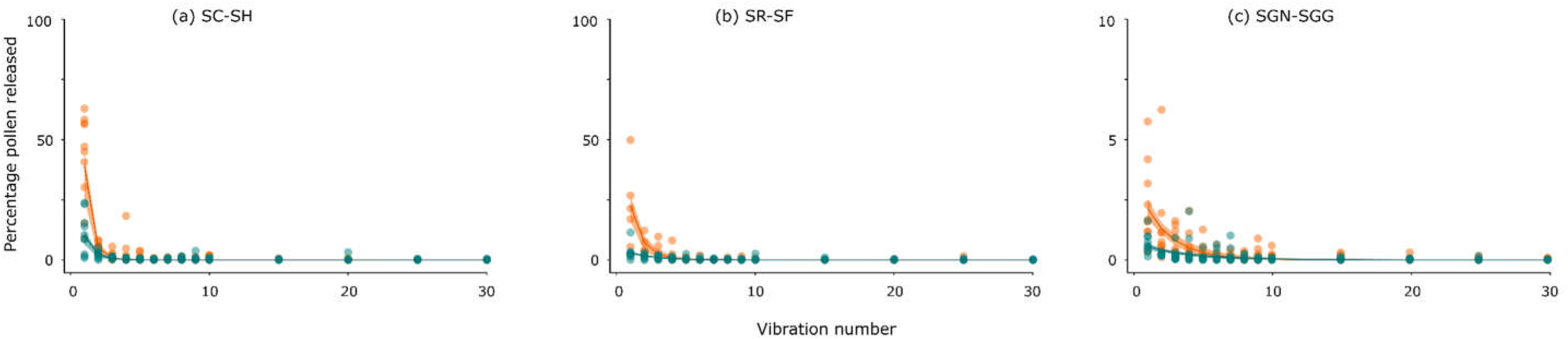
Pollen dispensing curves for three pairs of taxa in *Solanum* sect. *Androceras*, belonging to thee phylogenetic clades: (a) Series *Violaceiflorum* (b) Series *Androceras*, and (c) Series *Pacificum*. Each point shows the percentage of pollen released per vibration. Orange curves show the small flower type, and blue curves show the large flower type. These curves were fitted using *nls* in R and are based on the combined data of all flowers of a species. Confidence intervals (95%) were fitted using *predictNLS*. Species names as in Figure 1.

**Figure 3.**
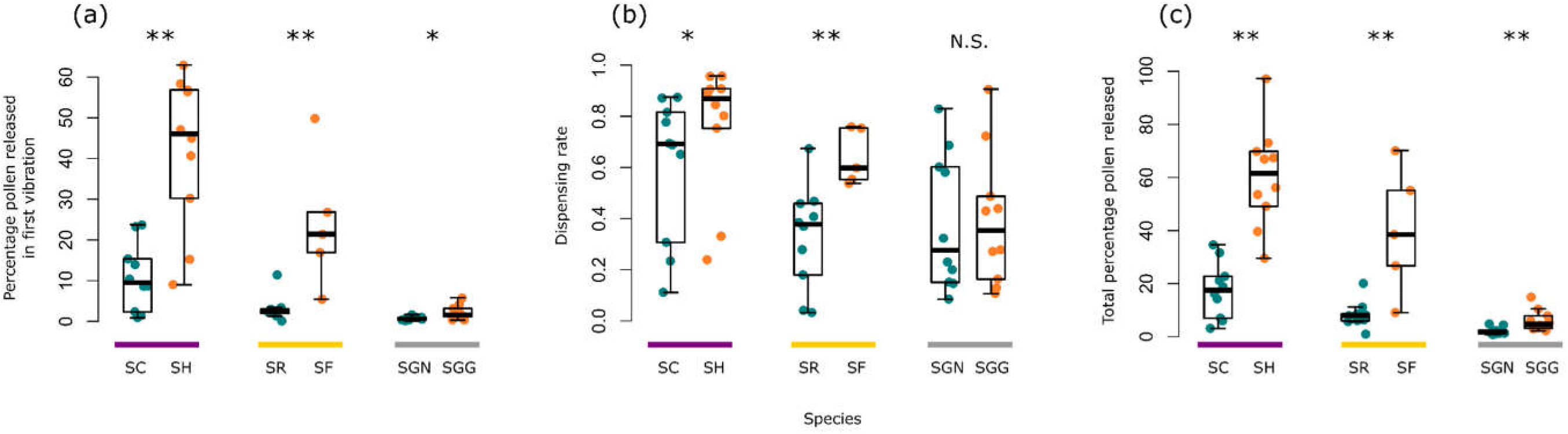
(a) The percentage pollen released in the first vibration is compared between flower types within the three clades, indicated by the purple, yellow and grey lines. For all clades, the small flower type released more pollen in the first vibration than the large flower type. (b) The dispensing rate is compared between flower types within the three clades. For two clades, the small flower type released pollen faster than the large flower type. (c) The total percentage pollen released across 30 vibrations is compared between flower types within clades. For all clades, the small flower type released more of its pollen than the large flower type. Orange curves show the small flower type, and blue curves show the large flower type. Species names as in Figure 1. * indicates 0.01 < *P* < 0.05; ** indicates 0.001 < *P* < 0.01.

### Variation in pollen dispensing schedules between flower types

The percentage of pollen released in the first vibration varied between flower types, and for all three phylogenetic clades, more pollen was released in the first vibration for small flower types than large flower types (SC-SH: W = 8, p < 0.001, p_adjusted_ = 0.002; SR-SF: W = 1, p = 0.001, p_adjusted_ = 0.003; SGN-SGG: W = 18, p = 0.01, p_adjusted_ = 0.01; Table 1; Figs. 2 & 3). Across our six taxa, *Solanum heterodoxum* released the largest percentage of pollen grains in the first vibration (46.03% ± 5.88; median ± se) and *S. grayi* var. *grandiflorum* released the smallest (0.58% ± 0.13; Table 1). For most species, the amount of pollen released in the first vibration represented more than half of the pollen that was released during the 30 vibrations (Table 2).

**Table 2.** Mixed effect models were used to assess the association between pollen dispensing schedules and various anther traits. In (a) a negative binomial mixed effect model was conducted with species identity as random factor. We used the number of pollen grains released in the first vibration as response variable, and tested whether this is associated with the anther wall area and anther pore area of the feeding and pollinating anthers separately. We used the total amount of pollen in a flower as offset in the model. In (b), we conducted a logistic mixed-effects model with species identity as random factor. We used untransformed pollen dispensing rates (*b*; ranging from 0 to 1) as response variable (with high values indicating most pollen is released in few vibrations) and tested whether this was associated with the anther wall area and anther pore area of the feeding and pollinating anthers separately. Significance at *P* < 0.05 is indicated in bold.

| **(a) Pollen released in first vibration** |  |  |  |
| --- | --- | --- | --- |
| Variable | Coefficient | SE | *P*-value |
| Intercept | -3.2191 | 0.8657 |  |
| *Feeding anther:* |  |  |  |
| Area (mm<sup>2</sup>) | 0.0299 | 0.0319 | 0.35 |
| Pore size (mm<sup>2</sup>) | 2.1685 | 1.7722 | 0.22 |
| *Pollinating anther:* |  |  |  |
| Area (mm<sup>2</sup>) | **-0.0751** | **0.0336** | **0.03** |
| Pore size (mm<sup>2</sup>) | 5.4196 | 4.3688 | 0.21 |
| **(b) Pollen dispensing rate (*b*)** |  |  |  |
| Fixed effect | Coefficient | SE | *P*-value |
| Intercept | -0.2263 | 1.2193 |  |
| *Feeding anther:* |  |  |  |
| area | 0.0471 | 0.0313 | 0.13 |
| pore size | 0.3294 | 4.0790 | 0.94 |
| *Pollinating anther:* |  |  |  |
| area | **-0.0973** | **0.0499** | **0.05** |
| pore size | 8.8027 | 16.7648 | 0.60 |

For two of the three clades, the dispensing rates were higher in small flowered taxa than large flowered taxa, indicating differences in the shapes of the curves. Specifically, the dispensing rates for small flowered taxa were higher than those of large flower types for the SR-SF clade (W = 3, p = 0.005, p_adjusted_ = 0.01) and marginally significant for the SC-SH clade (W = 24, p = 0.05, p_adjusted_ = 0.10), but not for the SGN-SGG clade (W = 48, p = 0.91, p_adjusted_ = 0.91; Figs. 2 & 3; Table 2). The fast dispensing rates in small flowered taxa show that these taxa required fewer vibrations to release all of the available pollen than large flowered taxa.

None of the six taxa released all of their pollen during 30 vibrations, despite no pollen being released after 30 vibrations. The total amount of pollen released varied strongly between the species of each phylogenetically independent contrast, with large flowered taxa releasing proportionally fewer pollen grains in 30 vibrations than their small flowered sister taxon (SC-SH: W = 2, p < 0.001, p_adjusted_ < 0.001; SR-SF: W = 48, p = 0.002, p_adjusted_ = 0.005; SGN-SGG: W = 88, p = 0.002, p_adjusted_ = 0.005; Fig. 3; Table 1). We also observed differences between clades, where the largest proportion of pollen was released by the SC-SH clade and the smallest proportion was released by the SGG-SGN clade.

### Effect of vibration properties on pollen dispensing schedules

The application of different peak vibration velocities resulted in different pollen dispensing schedules. Generally, lower vibration velocities resulted in slower pollen release and less total pollen released over 30 vibrations.

We tested whether the percentage of pollen released in the first vibration varied with vibration velocity, and we found effects of vibration velocity for both the large flowered *S. citrullifolium* (F_2,27_ = 11.44, p < 0.001) and the small flowered *S. heterodoxum* (F_2,25_ = 21.86, p < 0.001) (Figs. 4 & 5). Tukey posthoc tests showed that for *S. citrullifolium*, a velocity of 80 mm/s released more pollen in the first vibration than both lower velocity vibrations (SC_40mm/s_: p = 0.002; SC_20mm/s_: p < 0.001). For each sequential lower velocity vibration, we observed a more than four times decrease in the percentage pollen released (Table 1). Similarly, a velocity of 80 mm/s released more pollen in *S. heterodoxum* plants than both lower velocity vibrations (SH_40mm/s_: p < 0.001; SH_20mm/s_: p < 0.001), and we observed a nine times decrease in percentage pollen release between velocities of 80 mm/s and 40 mm/s. No differences in pollen released in the first vibration were present between velocities of 40 mm/s and 20 mm/s for either species (SC: p = 0.85; SH: p = 0.76).

**Figure 4.**
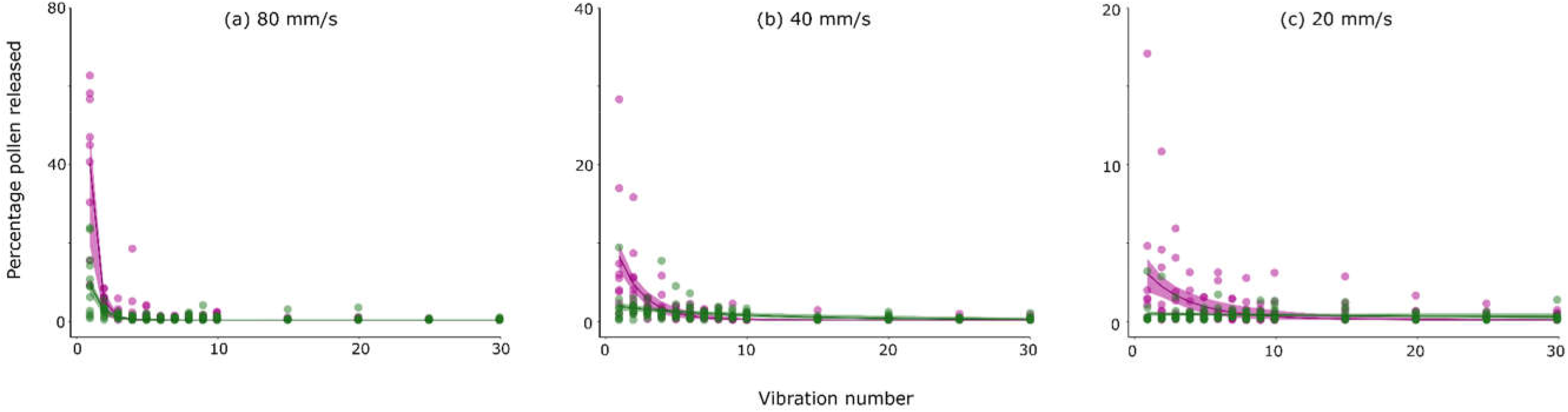
Pollen dispensing curves for *Solanum citrullifolium* (in green; large flower type) and *S. heterodoxum* (in pink; small flower type) when (a) 80 mm/s, (b) 40 mm/s, and (c) 20 mm/s vibration velocities were applied to flowers. Each point shows the percentage of pollen released per vibration. These curves were fitted using *nls* in R and are based on the combined data of all flowers of a species. Confidence intervals (95%) were fitted using *predictNLS*.

Further, we found differences in the pollen dispensing rates when different vibration velocities were applied. These differences were observed both in *S. citrullifolium* (F_2,27_ = 10.64, p < 0.001) and in *S. heterodoxum* (F_2,25_ = 11.50, p < 0.001) (Fig. 5). For *S. citrullifolium*, a velocity of 80 mm/s released pollen quicker than velocities of 40 mm/s (p = 0.001) and 20 mm/s (p < 0.001), which shows that more vibrations are required to release the available pollen grains when vibration velocities are lower. In particular, we observed a three-fold decrease in dispensing rate with a decrease in vibration velocity. Similarly, a velocity of 80 mm/s released pollen quicker in *S. heterodoxum* than velocities of 40 mm/s (p = 0.02) and 20 mm/s (p < 0.001). However, no differences in pollen dispensing rates were detected between velocities of 40 mm/s and 20 mm/s (SC: p = 0.91; SH: p = 0.45).

**Figure 5.**
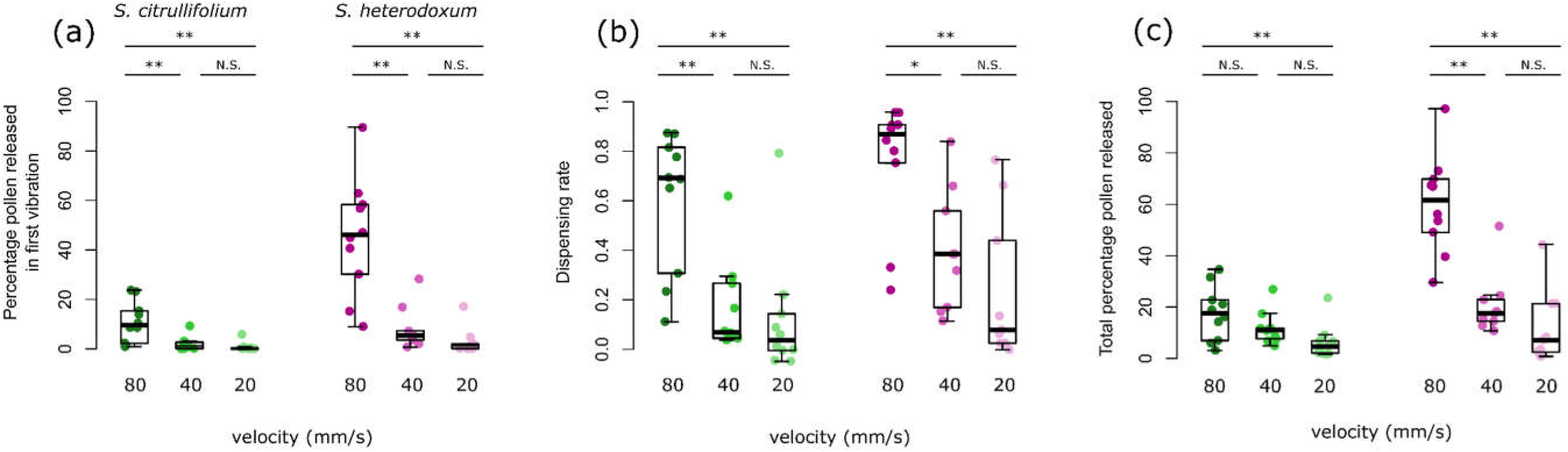
The percentage pollen released in the first vibration (a), the dispensing rate (b), and the total percentage pollen released across 30 vibrations (c) are compared between vibration velocities for *S. citrullifolium* (in green; large flower type) and *S. heterodoxum* (in pink; small flower type). * indicates 0.01 < p < 0.05; ** indicates 0.001 < p < 0.01.

In addition to changes in the shape of the pollen dispensing curves, differences in vibration velocity also influenced the total amount of pollen grains that were ejected, both for *S. citrullifolium* (F_2,27_ = 4.906, p = 0.01, Figs. 5 & 6) and for *S. heterodoxum* (F_2,25_ = 25.67, p < 0.001, Figs. 5 & 6). For *S. citrullifolium*, Tukey posthoc tests showed a significant decrease in the total amount of pollen released between 80 and 20 mm/s (p = 0.01), but not between 80 and 40 mm/s (p = 0.26) nor between 40 and 20 mm/s (p = 0.28). Similarly, Tukey posthoc tests showed a significant decrease in the total amount of pollen released for *S. heterodoxum* between 80 and 40 mm/s (p < 0.001) and between 80 and 20 mm/s (p < 0.001), but not between 40 and 20 mm/s (p = 0.41).

### Association between pollen dispensing schedules and anther traits

We tested whether anther traits (i.e., anther wall area and anther pore size) of the feeding and pollinating anthers relate to pollen dispensing, as hypothesized by Harder & Barclay (1978). The amount of pollen released in the first vibration was negatively associated with the pollinating anther wall area (z = −2.232, p = 0.03, Table 2), showing that flowers with larger pollinating anthers release less pollen in the first vibration than those with smaller pollinating anthers. Similarly, pollen dispensing rates were negatively associated with the pollinating anther wall area (z = −1.952, p = 0.05, Table 2), showing that flowers with larger pollinating anthers release pollen more slowly than flowers with smaller anthers.

## Discussion

Our study is the first to systematically investigate the effects of anther morphology and vibration properties on pollen dispensing schedules in buzz-pollinated plants. When applying bee-like vibrations directly to anthers, we found consistent differences in pollen release rates between anther types, where the large anther type released pollen more gradually than the small anther type. Higher vibration velocities resulted in more pollen released in the first vibration and faster pollen release rates, irrespective of the anther type. We thus show that both anther morphology and bee vibrations are associated with pollen release schedules, and likely pollen export, in buzz-pollinated plants. Additionally, we found that larger pollinating anther areas were associated with less pollen released in the first vibration and a slower dispensing rate, showing the potential for relating individual anther traits to pollen release in poricidal taxa.

### Pollen dispensing schedules in buzz-pollinated taxa

Many plant species stagger the pollen release of individual flowers across hours or days (Harder & Barclay, 1994; Sargent, 2003; Castellanos *et al.*, 2006b; Li *et al.*, 2014; Dellinger *et al.*, 2019c), and here we show that flowers can also limit pollen release across shorter timeframes (i.e., seconds and minutes). For all six taxa, pollen was released across multiple vibrations, with a smaller amount of pollen released in each successive vibration. This suggests that multiple visits or extended visits by vibrating pollinators are required to extract all of the pollen available for release (see Larson & Barrett, 1999), and that bees would receive the largest pollen rewards in the first vibration. Most flowers required between three and ten vibrations to release all available pollen grains, which translates to 6 to 20 seconds of buzzing. Our results align with the field observations made by Bowers (1975) which showed that bees tend to spend more time (i.e., 3 – 15 seconds) on newly opened *S. rostratum* flowers and less time on previously visited flowers, presumably matching their visitation lengths with pollen availability.

In line with work on non-poricidal taxa, we also found that pollen release is staggered over longer timeframes. Four of our six taxa released only a small percentage of their total pollen across the thirty vibrations we applied (<20%, median, Fig. 3), even though no pollen was released during the last few vibrations of each trial (see Fig. 2). Because this secondary pollen dosing mechanism does not seem to result from anther morphology, it is likely that pollen maturation or drying of pollen kitt is staggered in these plants, as is seen in other buzz-pollinated taxa (Corbet *et al.*, 1988; King & Ferguson, 1994; King & Buchmann, 1996). Staggering pollen maturation can also allow dynamic adjustment of pollen release schedules to pollinator visitation rates. For instance, if a flower that staggers pollen maturation receives its first visit on the second day of anthesis (e.g., under low visitation rates), then a larger quantity of pollen will be available for extraction than a flower which is visited on the first day (Harder & Barclay 1994). This dynamic adjustment of pollen release to pollinator visitation rates is predicted by theoretical models (Harder & Wilson 1994) and has been shown to occur in other poricidal taxa (Harder & Barclay, 1994; Dellinger *et al.*, 2019c).

### Shifts in pollen dispensing schedules between flower types

Theoretical models predict that pollen release schedules should be optimized to pollinator grooming behaviour and pollinator visitation rates (Harder & Thomson, 1989; Harder & Wilson, 1994; LeBuhn & Holsinger, 1998), and empirical work has shown support for these models (Sargent, 2003; Castellanos *et al.*, 2006a; Li *et al.*, 2014). A notable study by Castellanos *et al*. (2006) showed that parallel shifts in *Penstemon* between bee and hummingbird pollination are associated with shifts in anther morphology and pollen dispensing schedules. Similarly, we show that closely-related species in *Solanum* sect. *Androceras*, that have undergone parallel shifts in anther morphology (Vallejo-Marín *et al.*, 2014), have also undergone parallel shifts in their pollen dispensing schedules. If these pollen dispensing schedules are adaptations to flower visitors, then the quick release of pollen by the small flowered *Solanum* species is indicative of adaptation to a pollination strategy that results in high per-visit pollen transfer efficiency, whereas plants in the large flower group are likely frequently visited by pollinators that groom large quantities of pollen. Importantly, our work shows that these changes in pollen dispensing schedules can occur within a pollination syndrome.

Several non-mutually exclusive drivers have been proposed for the parallel shifts in morphology within *Solanum* sect. *Androceras*, and these relate well to the differences in pollen dispensing schedules we observe. Firstly, previous work has suggested that the small flowered taxa in this clade exhibit traits that correspond to self-pollination (Vallejo-Marín *et al.*, 2014; Rubini-Pisano *et al.*, in prep). The rapid release of pollen in small flowered taxa would correspond to self-pollination where plants do not need to mitigate the costs associated with pollinator grooming. Alternately, Whalen (1978) suggested that the size of flowers in this clade resulted from character displacement, where the size of pollinators matches the size of flowers. Whalen (1978) observed small bees visiting the small flower types, and the quick pollen release rates might reflect optimization to the visitation rates or grooming behaviours of these smaller solitary bees. However, pollination efficiently of these pollinator groups have not been assessed. He also observed larger bees, primarily bumblebees, pollinating the large flowers (Whalen, 1978), and their frequent grooming or high visitation rates could be driving the slower pollen dispensing schedules. Finally, selection by smaller bees for smaller overall flower size and smaller anthers could indirectly affect the pollen dispensing rate, particularly as we found that smaller anther wall areas are associated with faster dispensing rates. Despite this possibility, the association between pollen dispensing schedules and anther traits that we demonstrate will result in different pollen export dynamics between anther types. Further, our results also show parallel shifts in secondary pollen dosing rates, which suggests that pollen dispensing schedules are not primarily by-products of selection on other phenotypic traits.

### The influence of vibration velocity on pollen dispensing schedules

For a variety of buzz-pollinated taxa, the amount of pollen released in a single vibration increases with vibration velocity when artificial vibrations are applied (Harder & Barclay, 1994; King & Buchmann, 1996; De Luca *et al.* 2013) as predicted by Buchmann and Hurley’s (1978) model. Here, we show for the first time that vibration velocity also influences the amount of pollen released in successive vibrations. Low velocity vibrations resulted in slower pollen dispensing rates, with less pollen dispensed during 30 vibrations. Accordingly, bees that produce low velocity vibrations are unlikely to extract large pollen quantities. These bees would either need to visit multiple flowers or they would potentially be discouraged from visiting these plants (Harder, 1990b; Nicholls & Hempel de Ibarra, 2017). Poricidal anthers could thus act as filter to insect taxa that cannot produce the necessary vibrations (De Luca & Vallejo-Marín, 2013; Sun & Rychtář, 2015), and restrict access to pollen rewards in a similar way in which long nectar tubes exclude insects with short probosces from accessing nectar rewards (Newman *et al.*, 2014; Santamaría & Rodríguez-Gironés, 2015; Zung *et al.*, 2015). This vibration filter might be particularly effective in discouraging visitation from small bees that produce lower velocity vibrations (De Luca *et al.*, 2013, 2019) and do not make contact with reproductive organs (Solís-Montero & Vallejo-Marín, 2017). However, some bee taxa might be able to adjust their vibration velocities or duration (Harder & Barclay, 1994; Morgan *et al.*, 2016; Russell *et al.*, 2016), resulting in increased pollen extraction at a higher energy cost to the insect. Field observations of the large flowered *S. rostratum* have shown that small buzzing bees frequently visit flowers (Solís-Montero *et al.*, 2015), but larger taxa that produce higher velocity vibrations (De Luca *et al.*, 2019), such as *Bombus* and *Xylocopa*, are common visitors (Whalen, 1978), as well as efficient pollinators (Solís-Montero & Vallejo-Marín, 2017).

Although bees that produce high velocity vibrations are the most effective at extracting pollen, attracting such visitors might not always be the best strategy for plants. Optimal dispensing schedules are contingent on pollinator grooming behaviours (Harder & Thomson, 1989), and if bees groom large amounts of pollen per visit, then slower dispensing (as induced by lower velocity vibrations) should theoretically result in higher plant fitness. Additionally, optimal dispensing schedules are dependent on pollinator visitation rates, and these are likely to be dependent on bee and plant community composition. Work by Sargent (2003) showed that temporal changes in pollinator community composition and visitation rates were associated with temporal within-species changes in pollen dispensing rates. Thus, if pollinator visitation rates are low, quick dispensing will be favoured, even if bees collect large amounts of pollen per visit. Optimal dispensing schedules are thus expected to be adapted to the ecological community context, as well as the behaviour of bees.

### Anther traits and pollen dispensing schedules

Our results clearly show that pollen dispensing schedules are associated with both anther type and vibration properties, yet connecting pollen release schedule variation to specific anther traits is much more challenging. We find that larger anther areas are associated with less pollen released in the first vibration. This effect is statistically significant for pollinating anthers but not for feeding anthers (Table 2a), suggesting that other aspects of the morphology and material properties of anthers might be important. Similarly, we find that larger anther areas are associated with reduced rates of pollen release (not significant in feeding anthers; Table 2b). Our results contrast the expectation generated by the Buchmann & Hurley (1978) model that large anther locule areas should be associated with higher dispensing rates. However, their model also predicts a negative association between anther locule volume and pollen dispensing rates. Because anther wall area and anther locule volume are linked, we are potentially detecting the effects of locule volume rather than locule wall area. Quantifying the internal volume of anther locules is difficult but possible using techniques such as X-ray micro-computer tomography scanning (Dellinger *et al*., 2019a), and our work suggests that this is likely an important variable to measure. The lack of a statistically significant association between pore size and pollen release parameters is intriguing, but perhaps not unexpected given smaller variation in pore sizes than anther areas (see Fig. 1), as well as the conflicting effects of pore area on pollen release in the theoretical model of Buchmann and Hurley.

Our results highlight the need for more empirical studies of pollen dispensing schedules in buzz-pollinated plants, as well as for developing biophysical models of buzz-pollination that incorporate additional aspects of the morphology, geometry and material properties of flowers. The vibrations that the anthers experience depend both on the characteristics of bee vibrations (Switzer *et al.*, 2019; Pritchard & Vallejo-Marín, 2020) and on the properties of the floral structures. For example, the transmission of vibrations through the flower can alter the vibration that the anther experiences, e.g., by dampening the vibration velocity (King, 1993; Arroyo-Correa *et al.*, 2019), which can vary between closely-related taxa (Arroyo-Correa *et al.*, 2019). In contrast, if the bee vibrates the anther at its natural frequency (Nunes *et al*., 2020), resonance could amplify anther velocity and result in higher pollen removal (King & Buchmann, 1996). Joint modelling and experimental approaches to buzz pollination have the potential to help us understand the biomechanics and function of a fascinating biological interaction involving thousands of plant and bee species (Buchmann 1983; Cardinal *et al.*, 2018).

## Acknowledgments

We thank the members of the Vallejo-Marin lab for fruitful conversations on buzz-pollination and help with plant maintenance, with particular thanks to Carlos Pereira Nunes. We thank Ian Washbourne and George MacLeod for assistance with the use of the particle counter and SEM respectively, and Boris Igic and Lislie Solís-Montero for help with fieldwork to collect seed material. The project was made possible by the Royal Society of London and the Newton Fund through a Newton International Fellowship to JEK (NIF/R1/181685), by a Scottish Plant Health License (PH/38/2018-2020), and a research grant from The Leverhulme Trust (RPG-2018-235) to MVM.

## References

Arceo-Gómez G, Martínez ML, Parra-Tabla V, García-Franco JG. 2011. Anther and stigma morphology in mirror-image flowers of *Chamaecrista chamaecristoides* (Fabaceae): Implications for buzz pollination. Plant Biology 13: 19–24.

Arroyo-Correa B, Beattie C, Vallejo-Marín M. 2019. Bee and floral traits affect the characteristics of the vibrations experienced by flowers during buzz pollination. Journal of Experimental Biology 222:jeb198176.

Bowers KAW. 1975. The pollination ecology of *Solanum rostratum* (Solanaceae). American Journal of Botany 62: 633–638.

Buchmann SL. 1983. Buzz pollination in angiosperms. *Handbook of Experimental Pollination Biology*: In: Jones CE, Little RJ, eds. Handbook of experimental pollination biology. New York, NY: Scientific and Academic Editions, 73–113.

Buchmann SL, Hurley JP. 1978. A biophysical model for buzz pollination in angiosperms. Journal of Theoretical Biology 72: 639–657.

Cardinal S, Buchmann SL, Russell AL. 2018. The evolution of floral sonication, a pollen foraging behavior used by bees (Anthophila). Evolution 72: 590–600.

Castellanos MC, Wilson P, Keller SJ, Wolfe AD, Thomson JD. 2006. Anther evolution: pollen presentation strategies when pollinators differ. The American Naturalist 167: 288–296.

Corbet SA, Chapman H, Saville N. 1988. Vibratory pollen collection and flower form: bumble-bees on *Actinidia*, *Symphytum*, *Borago* and *Polygonatum*. Functional Ecology 2: 147.

Corbet SA, Huang SQ. 2014. Buzz pollination in eight bumblebee-pollinated *Pedicularis* species: Does it involve vibration-induced triboelectric charging of pollen grains? Annals of Botany 114: 1665–1674.

De Luca PA, Buchmann S, Galen C, Mason AC, Vallejo-Marín M. 2019. Does body size predict the buzz-pollination frequencies used by bees? Ecology and Evolution 9: 4875–4887.

De Luca PA, Bussière LF, Souto-Vilaros D, Goulson D, Mason AC, Vallejo-Marín M. 2013. Variability in bumblebee pollination buzzes affects the quantity of pollen released from flowers. Oecologia 172: 805–816.

De Luca PA, Vallejo-Marín M. 2013. What’s the ‘buzz’ about? The ecology and evolutionary significance of buzz-pollination. Current Opinion in Plant Biology 16: 429–435.

Dellinger AS, Artuso S, Pamperl S, Michelangeli FA, Penneys DS, Fernández-Fernández DM, Alvear M, Almeda F, Scott Armbruster W, Staeder Y,et al. 2019a. Modularity increases rate of floral evolution and adaptive success for functionally specialized pollination systems. Communications Biology 2: 453.

Dellinger AS, Chartier M, Fernández-Fernández D, Penneys DS, Alvear M, Almeda F, Michelangeli FA, Staedler Y, Armbruster WS, Schonenberger J. 2019b. Beyond buzz-pollination – departures from an adaptive plateau lead to new pollination syndromes. New Phytologist 221: 1136–1149.

Dellinger AS, Pöllabauer L, Loreti M, Czurda J, Schönenberger J. 2019c. Testing functional hypotheses on poricidal anther dehiscence and heteranthery in buzz-pollinated flowers. Acta ZooBot Austria 156: 197–214.

Dulberger R, Smith MB, Bawa KS. 1994. The stigmatic orifice in *Cassia*, *Senna*, and *Chamaecrista* (Caesalpiniaceae): Morphological variation, function during pollination, and possible adaptive significance. American Journal of Botany 81: 1390–1396.

Faegri K. 1986. The solanoid flower. Transactions of the Botanical Society of Edinburgh 45: 51–59.

Fick SE, Hijmans RJ. 2017.WorldClim 2: new 1km spatial resolution climate surfaces for global land areas. International Journal of Climatology 37(12): 4302–4315.

Harder LD. 1990a. Behavioral responses by bumble bees to variation in pollen availability. Oecologia 85: 41–47.

Harder LD. 1990b. Behavioral responses by bumble bees to variation in pollen availability. Oecologia 85: 41–47.

Harder LD, Barclay RMR. 1994. The functional significance of poricidal anthers and buzz pollination: controlled pollen removal from *Dodecatheon*. Functional Ecology 8: 509–517.

Harder LD, Johnson SD. 2009. Darwin’ s beautiful contrivances: evolutionary and functional evidence. New Phytologist 183: 530–545.

Harder LD, Thomson JD. 1989. Evolutionary options for maximizing pollen dispersal of animal-pollinated plants. The American Naturalist 133: 323–344.

Harder LD, Wilson WG. 1994. Floral evolution and male reproductive success: Optimal dispensing schedules for pollen dispersal by animal-pollinated plants. Evolutionary Ecology 8: 542–559.

Harris JA. 1905. The dehiscence of anthers by apical pores. Missouri Botanical Garden Report 1905: 167–257.

Holsinger KE, Thomson JD. 1994. Pollen discounting in *Erythronium grandiflorum*: mass-action estimates from pollen transfer dynamics. The American Naturalist 144: 799–812.

King MJ. 1993. Buzz foraging mechanism of bumble bees. Journal of Apicultural Research 32: 41–49.

King MJ, Buchmann SL. 2003. Floral sonication by bees: Mesosomal vibration by *Bombus* and *Xylocopa*, but not *Apis* (Hymenoptera: Apidae), ejects pollen from poricidal anthers. Journal of Kansas Entomological Society 76: 295–305.

King MJ, Buchmann SL, Spangler H. 1996. Activity of asynchronous flight muscle from two bee families during sonication (buzzing). Journal of Experimental Biology 199: 2317–2321.

King MJ, Ferguson AM. 1994.Vibratory collection of *Actinidia deliciosa* (kiwifruit) pollen. Annals of Botany 74: 479–482.

Knapp S. 2002. Floral diversity and evolution in the Solanaceae. In: Developmental genetics and plant evolution. 267–297.

Larson BMH, Barrett SCH. 1999.The pollination ecology of buzz-pollinated *Rhexia virginica* (Melastomataceae). American Journal of Botany 86: 502–511.

LeBuhn G, Holsinger K. 1998. A sensitivity analysis of pollen-dispensing schedules. Evolutionary Ecology 12: 111–121.

Li XX, Wang H, Gituru RW, Guo YH, Yang CF. 2014. Pollen packaging and dispensing: Adaption of patterns of anther dehiscence and flowering traits to pollination in three *Epimedium* species. Plant Biology 16: 227–233.

Minnaar C, Anderson B, Jager ML De, Karron JD. 2019. Plant-pollinator interactions along the pathway to paternity. Annals of Botany 123: 225–245.

Morgan T, Whitehorn P, Lye GC, Vallejo-Marín M. 2016. Floral sonication is an innate behaviour in bumblebees that can be fine-tuned with experience in manipulating flowers. Journal of Insect Behavior 29: 233–241.

Newman E, Manning J, Anderson B. 2014. Matching floral and pollinator traits through guild convergence and pollinator ecotype formation. Annals of Botany 113: 373–384.

Nicholls E, Hempel de Ibarra N. 2017. Assessment of pollen rewards by foraging bees. Functional Ecology 31: 76–87.

Nunes CEP, Nevard L, Montealegre-Zapata F, Vallejo-Marin M. 2020. Are flowers tuned to buzzing pollinators? Variation in the natural frequency of stamens with different morphologies and its relationship to bee vibrations. bioRxiv. doi: https://doi.org/10.1101/2020.05.19.104422

Ollerton J, Winfree R, Tarrant S. 2011. How many flowering plants are pollinated by animals? Oikos 120: 321–326.

Pritchard DJ, Vallejo-Marín M. 2020. Floral vibrations by buzz-pollinating bees achieve higher frequency, velocity and acceleration than flight and defence vibrations. Journal of Experimental Biology. jeb.220541 doi: 10.1242/jeb.220541

R Core Team. 2019. R: A language and environment for statistical computing. R Core Team. Vienna, Austria. http://www.R-project.org/

Rosi-Denadai CA, Araújo PCS, Campos LA de O, Cosme L, Guedes RNC. 2018. Buzz-pollination in Neotropical bees: genus-dependent frequencies and lack of optimal frequency for pollen release. Insect Science 27: 133–142.

Rubini Pisano A, Vallejo-Marín M, Benitez-Vieyra S, Fornoni J. In review. From large to small flowers in *Solanum* (Section *Androceras*): a multivariate perspective. Annals of Botany.

Russell AL, Leonard AS, Gillette HD, Papaj DR. 2016. Concealed floral rewards and the role of experience in floral sonication by bees. Animal Behaviour 120: 83–91.

Santamaría L, Rodríguez-Gironés M a. 2015. Are flowers red in teeth and claw? Exploitation barriers and the antagonist nature of mutualisms. Evolutionary Ecology 29: 311–322.

Sargent RD. 2003. Seasonal changes in pollen-packaging schedules in the protandrous plant *Chamerion angustifolium*. Population Ecology 135: 221–226.

Särkinen T, Bohs L, Olmstead RG, Knapp S. 2013. A phylogenetic framework for evolutionary study of the nightshades (Solanaceae): A dated 1000-tip tree. BMC Evolutionary Biology 13: 214.

Schlindwein C, Wittmann D, Martins CF, Hamm A., Siqueira JA, Schiffler D, MacHado IC. 2005. Pollination of *Campanula rapunculus* L. (Campanulaceae): How much pollen flows into pollination and into reproduction of oligolectic pollinators? Plant Systematics and Evolution 250: 147–156.

Schneider CA, Rasband WS, Eliceiri KW. 2012. NIH Image to ImageJ: 25 years of image analysis. Nature Methods 9: 671–675.

Solis-Montero L, Vergara C, Vallejo-Marín M. 2015. High incidence of pollen theft in natural populations of a buzz-pollinated plant. Arthropod-Plant Interactions 9: 599–611.

Solis-Montero L, Vallejo-Marín M. 2017. Does the morphological fit between flowers and pollinators affect pollen deposition? An experimental test in a buzz-pollinated species with anther dimorphism. Ecology and Evolution 7: 2706–2715.

Stern SR, Weese T, Bohs LA. 2010. Phylogenetic relationships in *Solanum* section *Androceras* (Solanaceae). Systematic Botany 35: 885–893.

Switzer CM, Combes SA. 2017. Bumblebee sonication behavior changes with plant species and environmental conditions. Apidologie 48: 223–233.

Switzer C, Russell A, Papaj D, Combes S, Hopkins R. 2019. Sonicating bees demonstrate flexible pollen extraction without instrumental learning. Current Zoology 65:425–436.

Thomson JD. 1986. Pollen transport and deposition by bumble bees in *Erythronium*: influences of floral nectar and bee grooming. Journal of Ecology 74: 329–341.

Vallejo-Marín M, Manson JS, Thomson JD, Barrett SCH. 2009. Division of labour within flowers: Heteranthery, a floral strategy to reconcile contrasting pollen fates. Journal of Evolutionary Biology 22: 828–839.

Vallejo-Marín M, Silva EM Da, Sargent RD, Barrett SCH. 2010. Trait correlates and functional significance in flowering of heteranthery plants. 188: 418–425.

Vallejo-Marín M, Walker C, Friston-Reilly P, Solís-Montero L, Igic B. 2014. Recurrent modification of floral morphology in heterantherous Solanum reveals a parallel shift in reproductive strategy. Philosophical Transactions of the Royal Society B: Biological Sciences 369: 20130256.

Vallejo-Marín M. 2019. Buzz pollination: studying bee vibrations on flowers. New Phytologist 224: 1068–1074.

Whalen MD. 1978. Reproductive character displacement and floral diversity in *Solanum* section *Androceras*. Systematic Biology 3: 77–86.

Whalen MD. 1979. Allozyme variation and evolution in *Solanum* section *Androceras*. American Society of Plant Taxonomists 4: 203–222.

Zung JL, Forrest JRK, Castellanos MC, Thomson JD. 2015. Bee-to bird-pollination shifts in *Penstemon*: effects of floral-lip removal and corolla constriction on the preferences of free-foraging bumble bees. Evolutionary Ecology 29: 341–354.

