## Supporting information for "Pollen dispensing schedules in buzz-pollinated plants: Experimental comparison of species with contrasting floral morphologies"

Date of acceptance:

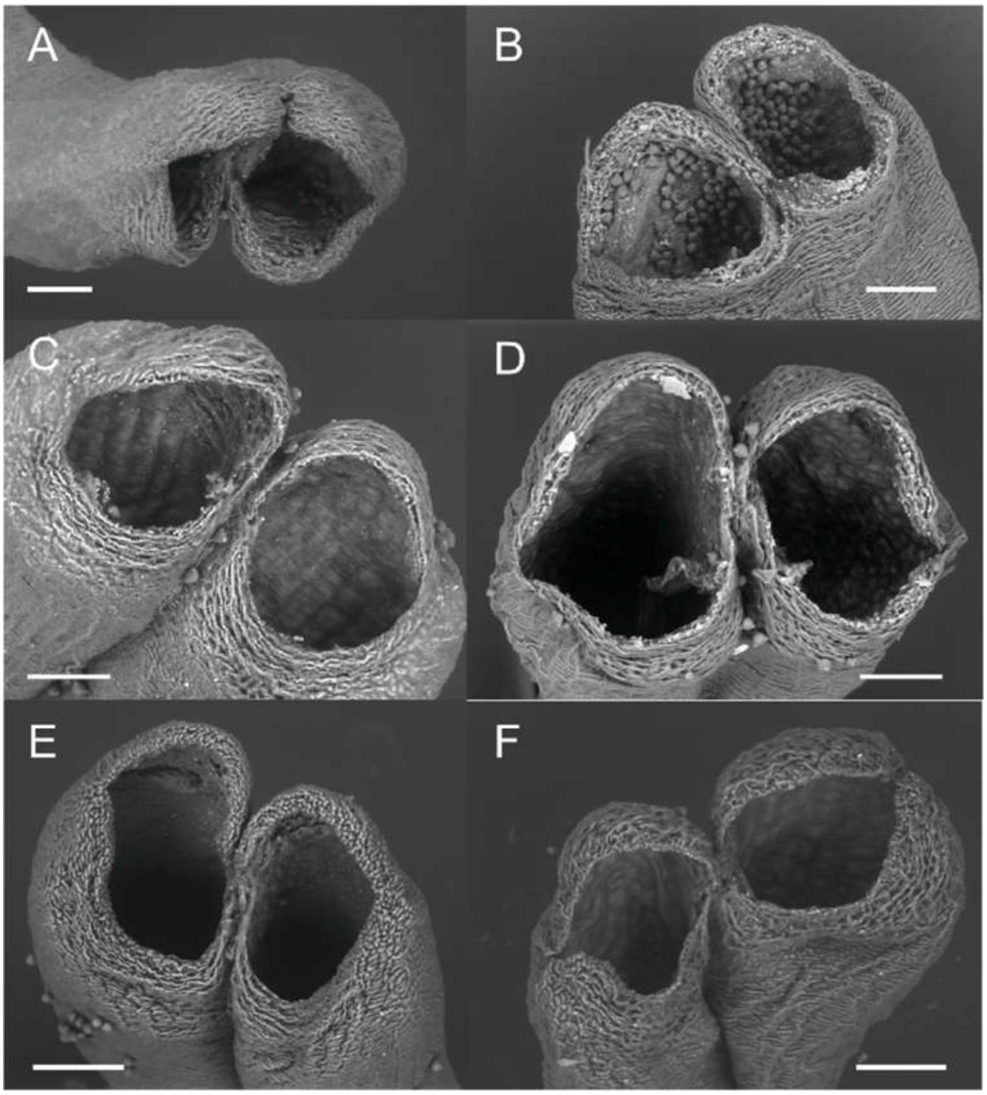


**Figure S1.** Pores of the pollinating anthers of (A) *Solanum rostratum*, (B) *S. fructu-tecto*, (C) *S. citrullifolium*, (D) *S. heterodoxum*, (E) *S. grayi* var. *grandiflorum*, and (F) *S. grayi* var. *grayi*. The panel on the left represent species of the ”large anther” type, and the panel on the right shows species of the “small anther” type.

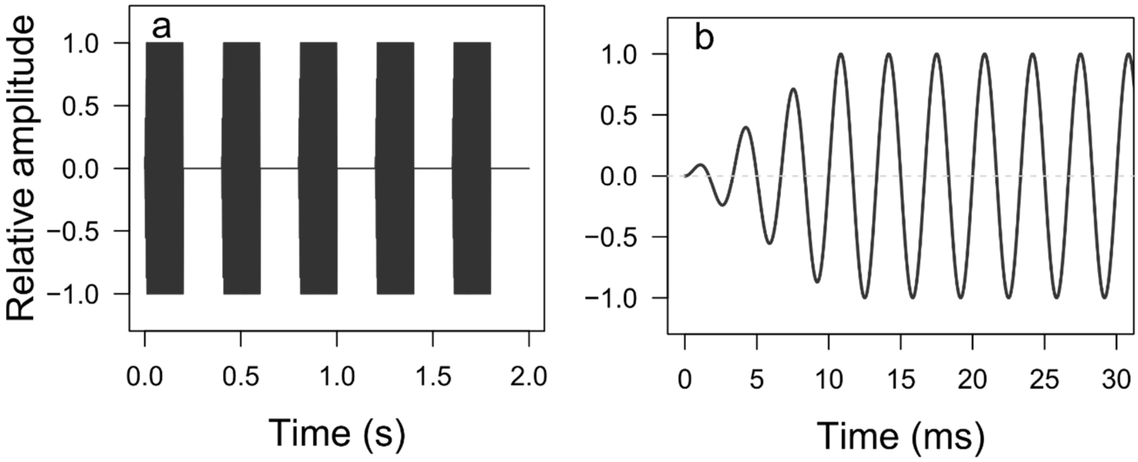


**Figure S2**. Artificial vibrations applied to anthers. (a) Each stimulus consisted of five short vibration pulses of 0.2 s long, with 0.2 s of silence between pulses. (b) The beginning of each 0.2 s pulse consisted of a short fade-in, which is similar to what bees produce and it ensures that the wave is transmitted in the expected manner.

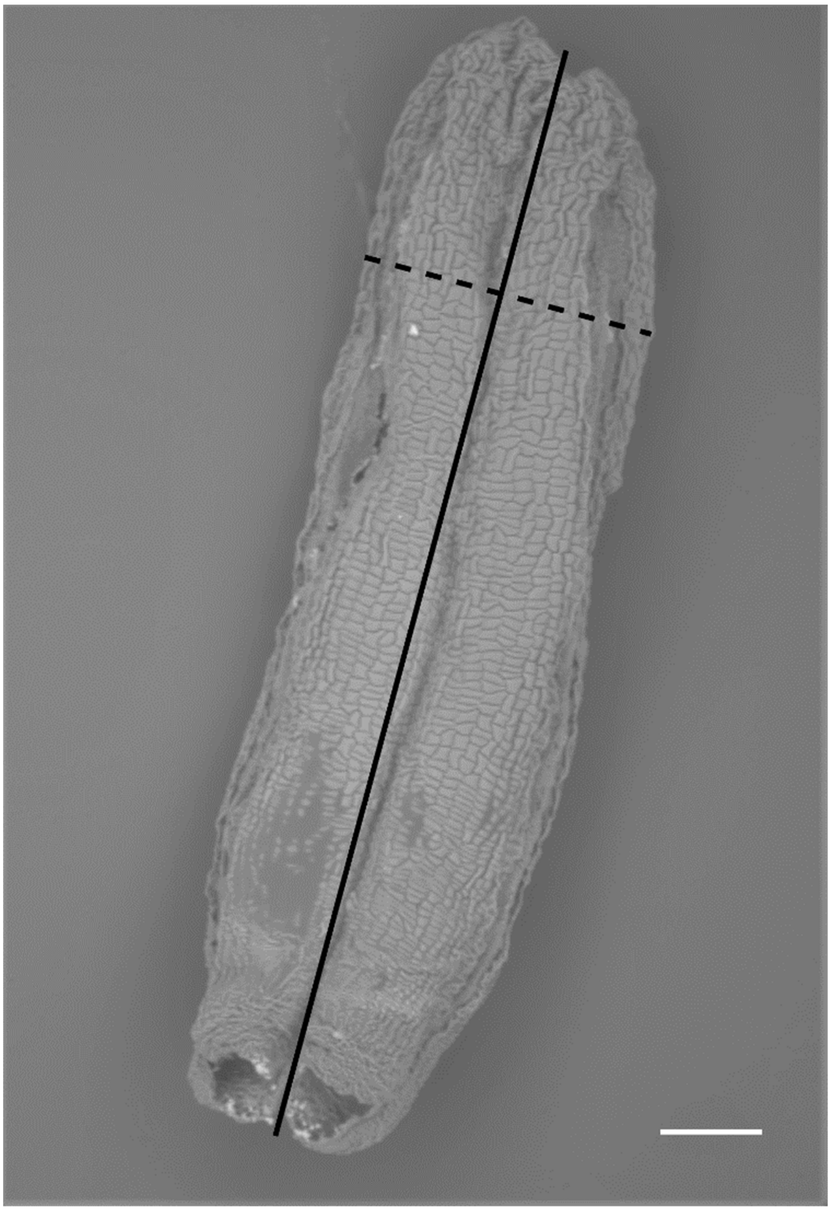

**Figure S3.** For each anther, the length (solid line) and breadth (dashed line) was measured. For each anther type within a flower the pore area was measure for one feeding and one pollinating anther. The photo shows a SEM image of a *Solanum fructu-tecto* feeding anther. The scale bar represents 200 µm.

**Table S1.** Plant material of *Solanum* section *Androceras* used in this study. The mean water vapour pressure (in kPa) was extracted from Worldclim2 using 1 $km^2$ spatial resolution. Data were extracted for each month separately and averaged to obtain the annual mean humidity for the sites where the seeds were originally collected.

| **Accession number** | **Species** | **Section** | **Population name** | **Latitude (N)** | **Longitude (W)** | **Mean humidity (in kPa)** |
|---|---|---|---|---|---|---|
| **199-7-3** | *S. citrullifolium* | Violaceiflorum | Nijmegen Collection | - | - | - |
| **11-PTEM-14, 15** | *S. heterodoxum* | | Teotihuacán, Estado de México | 19.68 | 98.84 | 1.08 |
| **10-s-81, 82, 86** | *S. rostratum* | Androceras | San Miguel de Allende, Querétaro | 20.90 | 100.45 | 1.00 |
| **10-AH-9, 24** | *S. fructu-tecto* | | Atitalaquia, Hidalgo | 20.06 | 99.21 | 1.15 |
| **08-s-78, 79** | *S. grayi* var. *grandiflorum* | Pacificum | Tejupilco, Estado de México | 18.85 | 100.13 | 1.59 |
| **07-s-194b, 195b, 196b** | *S. grayi* var. *grayi* | | Los Álamos, Sonora | 27.00 | 108.93 | 1.51 |
